# Region-specific differences and areal interactions underlying transitions in epileptiform activity

**DOI:** 10.1101/435156

**Authors:** Neela K. Codadu, R. Ryley Parrish, Andrew J. Trevelyan

## Abstract

Understanding the nature of epileptic state transitions remains a major goal for epilepsy research. Simple in vitro models offer unique experimental opportunities, which we exploit to show that such transitions can arise from shifts in the ictal source of the activity. These transitions reflect the fact that cortical territories differ both in the type of epileptiform activity they can sustain, and their susceptibility to drug manipulation. In the zero Mg^2+^ model, the earliest epileptiform activity is restricted to neocortical and entorhinal networks. Hippocampal bursting only starts much later, and triggers a marked transition in neo-/entorhinal cortical activity. Thereafter, the hippocampal activity acts as a pacemaker, entraining the other territories to their discharge pattern. This entrainment persists following transection of the major axonal pathways between hippocampus and cortex, indicating that it can be mediated through a non-synaptic route. Neuronal discharges are associated with large rises in extracellular [K^+^], but we show that these are very localised, and therefore are not the means of entraining distant cortical areas. We conclude instead that the entrainment occurs through weak field effects distant from the pacemaker, but which are highly effective at recruiting other brain territories that are already hyperexcitable. The hippocampal epileptiform activity appears unusually susceptible to drugs that impact on K^+^ conductances. These findings demonstrate that the local circuitry gives rise to stereotypical epileptic activity patterns, but these are also influenced by both synaptic and non-synaptic long-range effects. Our results have important implications for our understanding of epileptic propagation, and anti-epileptic drug action.

**Key points:**

- Local neocortical and hippocampal territories show different and sterotypical patterns of acutely evolving, epileptiform activity.
- Neocortical and entorhinal networks show tonic-clonic-like events, but the main hippocampal territories do not, unless it is relayed from the other areas.
- Transitions in the pattern of locally recorded epileptiform activity can be indicative of a shift in the source of pathological activity, and which may spread through both synaptic and non-synaptic means.
- Hippocampal epileptiform activity is promoted by 4-aminopyridine and inhibited by GABAB agonists, and appears far more sensitive to these drugs than neocortical activity.
- These signature features of local epileptiform activity can provide useful insight into the primary source of ictal activity, aiding both experimental and clinical investigation.

## Introduction

Epilepsy is a condition that is defined by sudden transitions, from a functional brain state into pathological states. These transitions are associated with dramatic changes also in the electrophysiological signals, and indeed, EEG recordings provide a very sensitive assay of brain states. The interpretation of these signals, though, is often difficult, and in most cases, we still do not understand what biological processes underlie the key shifts in the electrophysiological signal. They do, though offer great potential for providing advance warning about imminent seizures, and so warrant further study.

Epileptic transitions can arise from local network interactions (Bernard *et al.*, 2000; Ziburkus *et al.*, 2006; Huberfeld *et al.*, 2011; Trevelyan & Schevon, 2013; Avoli *et al.*, 2016), or cellular changes, such as intracellular chloride concentration (Dzhala *et al.*, 2010; Pavlov *et al.*, 2013; Ellender *et al.*, 2014; Pallud *et al.*, 2014). A role in these transitions has also been proposed for larger scale network interactions (Kramer *et al.*, 2012; Martinet *et al.*, 2017; Liou *et al.*, 2018). In these matters, it is extremely helpful to identify where is the source of the pathological discharge; this can then further provide insight into how it spreads. Most epileptic seizures are thought to arise from pathology located either in hippocampal, parahippocampal or neocortical circuits, but it remains unclear to what extent the pathological activity is set by the intrinsic excitability of the local networks (Traub & Wong, 1982; Miles & Wong, 1983; Prince & Connors, 1984; Dichter & Ayala, 1987; Ziburkus *et al.*, 2006) or interactions between the areas (Miles *et al.*, 1984; McCormick & Contreras, 2001). Brain slice preparations offer unique experimental opportunities both for recording, manipulating and isolating network activity. These preparations have yielded many insights into a wide range of topics from cellular excitability and synaptic interactions, up to network dynamics, for instance, by providing a framework to understand human recordings (Schevon *et al.*, 2012; Smith *et al.*, 2016) where the potential for invasive investigation is greatly limited.

We set out to investigate the role of interactions between brain areas in epileptic transitions. An important series of studies using the 0 Mg^2+^ model (Swartzwelder *et al.*, 1986b; Mody *et al.*, 1987; Anderson *et al.*, 1990; Dreier & Heinemann, 1990; 1991; Bragdon *et al.*, 1992; Morrisett *et al.*, 1993; Zhang *et al.*, 1995; Dreier *et al.*, 1998), characterised a notable transition, from early tonic-clonic patterns of epileptiform discharges, into a different, recurrent pattern of discharge. The nature of this critical transition, however, has remained a mystery. Of added interest is that this transition is associated with a marked change in the pharmaco-sensitivity of the pathological discharges (Heinemann *et al.*, 1994b). Since the various brain areas may differ in how epileptic discharges are manifest, we hypothesised that a key component of this transition might reflect changes in the level of involvement of different brain territories.

These prior studies all used brain slices prepared from adult rats, but have not been repeated using tissue from other species. We now show that the same evolution of activity is also seen in mouse brain slices, thereby opening up this phenomenon for further study in transgenic animals, carrying mutations relevant for human epilepsy. We then identify an important correlate of the transition, which is the surprisingly late involvement of hippocampal activation in this model, and which subsequently acts as a pacemaker, entraining activity in other cortical networks. Interestingly, the entrainment of overlying neocortex does not require intact synaptic pathways, but instead can arise from field effects secondary to focal discharges (Jefferys & Haas, 1982; Jefferys, 1995; Frohlich & McCormick, 2010; Anastassiou *et al.*, 2011). We further show that the entrainment does not happen through the diffusion of extruded K^+^, because the rise in extracellular [K^+^] associated with epileptiform discharges is very focal. Finally, the site of the dominant epileptiform activity in these preparations is highly sensitive to drugs that affect K^+^ conductance. The GABA_B_ agonist is a very powerful suppressor of the hippocampal focus, and shifts the slice back towards the neo- / entorhinal cortical pattern of tonic-clonic like discharges, whereas the K^+^ channel blocker, 4-aminopyridine strongly promotes hippocampal activity, far more rapidly than in the other areas, the exact opposite of the evolving pattern induced by 0 Mg^2+^. These results show that locally recorded transitions in the pattern of epileptiform discharge may arise from the new involvement of distally located epileptic circuits. These changes thus reflect which cortical territories are involved and how the activity spreads to other networks. These models illuminate a variety of epileptic phenomena, including the evolution of epileptic foci, sudden shifts from one foci to another, and how different cortical areas show distinctive patterns of epileptic discharge and propagation. As such, they can provide a wealth of metrics for comparing anti-epileptic drugs, and for understanding phenotypes in genetic models of epilepsy.

## Methods

### Ethical Approval

All animal handling and experimentation were done according to the guidelines laid by the UK Home Office and Animals (Scientific Procedures) Act 1986, and approved by the Newcastle University Animal Welfare and Ethical Review Body.

### Slice preparation

Young adult, male and female mice (wild-type C57-Bl6 strain, age 2-3 months) were sacrificed by schedule-1 method of cervical dislocation. The brains were removed and sliced horizontally (400μm thickness) in ice-cold artificial cerebrospinal fluid (ACSF; containing (in mM): 3, MgCl_2_; 126, NaCl; 26, NaHCO_3_; 3.5 KCl; 1.26 NaH_2_PO_4_; 10, glucose), using a Leica vibratome (Nussloch, Germany). Slices were immediately transferred to an interface tissue holding chamber and incubated for 1-2 hours at room temperature in ACSF containing (in mM): 2, CaCl_2_; 1MgCl_2_; 126 NaCl; 26 NaHCO_3_; 3.5 KCl; 1.26 NaH_2_PO_4_; 10, glucose). All the solutions were being bubbled continuously with carboxygen (95% O_2_ and 5% CO_2_).

### Electrophysiology

Slices were placed in an interface recording chamber and perfused with warmed ACSF (2-3mls/min, driven by a peristaltic pump (Watson-Marlow Pumps Limited, Cornwall UK; model 501U)). The temperature of the chamber and perfusate was maintained at 33-36°C using a closed circulating heater Grant FH16D (Grant instruments, Cambridge, UK). Extracellular field recordings were made using normal ACSF-filled 1-3MΩ borosilicate glass microelectrodes (GC120TF-10; Harvard apparatus, Kent) pulled using a Narishige electrode puller (PP-83, Narishige Scientific Instruments, Tokyo, Japan), and mounted headstages (10x DC pre-amp gain) held in Narishige YOU-1 micromanipulators. In experiments involving dissections, scalpel blades were used to make cuts in the slices after placing them in the recording chamber. Waveform signals were acquired using BMA-931 biopotential amplifier (Dataq instruments, Akron, USA), Micro 1401-3 ADC board (Cambridge Electronic Design, UK) and Spike2 version 7.10 software (Cambridge Electronic Design, UK). Signals were sampled at 10 kHz, amplified (gain: 200) and bandpass filtered (1-3000Hz). A CED4001-16 Mains Pulser (Cambridge Electronic Design, UK) was connected to the events input of CED micro 1401-3 ADC board and was used to remove 50Hz hum offline. Recordings were initiated while slices were still being perfused with normal ACSF, and only then was the perfusate switched to an epileptogenic ACSF solution lacking Mg^2+^ ions (0 Mg^2+^ACSF) or 100μM 4-aminopyridine (4AP).

Extracellular potassium (K^+^) was measured using single-barrelled K^+^-selective microelectrodes. The pipettes were pulled from nonfilamented borosilicate glass (Harvard Apparatus, Kent, UK), and the glass was exposed to vapour of dimethyl-trimethyl-silylamine (Sigma-Aldrich), baking at 2OO°C for 40 minutes, the pipettes were then backfilled with ACSF. A short column of the K^+^ sensor (Potassium ionophore I, cocktail B; Sigma-Aldrich, #99373) was taken into the tip of the salinized pipette by using slight suction. The recordings through the K^+^-sensor electrode was referenced to a second electrode filled with aCSF, and from the differential signal, we calculated the [K^+^]_o_ from calibration recordings made in an open bath, using sudden increments in [K^+^]_o_. We checked the stability of the electrodes at the start and end of each recording. Data from unstable electrode recordings were discarded. This provided a scaling factor S, of 55-59mV, where the K^+^ concentration at a given moment in time, t, was calculated from the differential voltage, V(t), as follows:

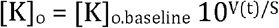

[K]_o.baseline_ for our experiments was 3.5mM.

### Data Analysis and Statistics

Data were analysed offline using Clampfit (Molecular Devices, CA, USA), Igor (WaveMetrics, Lake Oswego, OR) and Matlab R2015b (MathWorks, USA). The analysis of entrainment of epileptiform events was performed by deconvolution of an averaged “template” (Figure 5D) of electrophysiological discharges against the continuous trace from that same recording. In this way, the higher frequency components of these discharges are effectively removed. This is very helpful, because the cross-correlation analyses between hippocampal and neocortical discharges is optimal if the signal can be simplified essentially to the timing of the events, thereby minimizing any aliasing issues that might arise from these higher frequency components. The deconvolution was done by first creating a template of an average discharge (6-10 events), aligned by the time point at which they exceeded a threshold set at between 25-40% of the peak deflection. The templates were then used as a normalising filter on their respective raw traces, by deriving peak cross-correlation coefficients for the time-shifted template relative to the trace. This “template-filtered” trace (Figure 5D) removed most of the brain-region-specific fine structure of the individual discharges, but preserved their timing. Since the individual events in the late-stage activity are extremely reproducible, the peaks in this filtered trace tend towards 1. We used the cross-correlation between these template-filtered recordings as a measure of the entrainment of the two recording locations. Matlab code for these analyses is available from the authors upon request. A one-way analysis of variance (ANOVA), with a Tukey post hoc test was used for data with 3 or more groups. Groups of two were analyzed using the Student’s t-test. Significance was set at P ≤ 0.05 for all analyses. Multiunit activity was extracted from raw data by high-pass filtering it to >300Hz. Data is represented as mean ± s.e.m., and ‘n’ value is the number of brain slices, unless otherwise stated.

### Terminology

The terminology of epileptic discharges is problematic, reflecting the fact that there is a large range of activity patterns, and the equivalence of, or distinction between these is often hard to discern. This is particularly so for the term “interictal”, which in the clinical setting refers to electrophysiological activity which is clinically covert (if not completely so – see (Binnie *et al.*, 1987; Kleen *et al.*, 2010). These are typically rather short discharges, and consequently, animal researchers have taken this ephemeral nature to be the defining feature. Unfortunately when one examines the activity patterns closely, this term conflates two very different types of activity; an appreciation of the difference is critical for the understanding of this present study. The key distinction is the presence or absence of local intense activity, as defined by the presence of a significant high frequency component. The importance of this is that it helps distinguish between sites where there is local pathological activity, from those where the deviation in the recording reflects pathological activity that is elsewhere. Consequently, in this paper, we refer to the discharges during late stage activity as “spike and wave discharges”, a term that has been used previously to describe what appear to be comparable events. Readers should note however, that previous studies have described this activity pattern as “interictal”, but for the reasons outlined above, we prefer not to use this term.

## Results

### Region-specific patterns of evolving epileptiform activity

We investigated the evolution of epileptiform activity in horizontal brain slices, prepared from young adult (1-2 month old), wild-type C57B6J mice, following the removal of Mg^2+^ ions from the bathing medium (ACSF). Extracellular recordings were made at two or three locations, always including a hippocampal (CA1 or CA3) and a neocortical (temporal association areas) recording site, and in most slices, also recording from medial entorhinal cortex (Figure 1A). Following the washout of Mg^2+^ ions, there was a gradual build-up of epileptiform discharges, evolving in a highly characteristic way (Figure 1A). The earliest large field deflections in the raw traces were seen at all recording sites, although the events appeared far larger in the neocortex and entorhinal cortex. This early activity involved episodes of sustained rhythmic bursts suggestive of the temporal dynamics of clinical tonic-clonic discharges (Figure 1B). The mean number of tonic-clonic like events in the neocortical was 9.35 ± 0.73 per slice (range 4-17 events; n = 17 slices), before a second transition to regular epileptiform bursts (“late-stage activity pattern”), with individual bursts lasting a few hundred milliseconds, and occurring every 3.32 ± 0.38s (n = 10; Figure 2). This pattern of evolution has been described previously in rat brain slices (Swartzwelder *et al.*, 1986a; Mody *et al.*, 1987; Anderson *et al.*, 1990; Dreier & Heinemann, 1990; 1991; Bragdon *et al.*, 1992), but has been less studied in mice.

**Figure 1.**
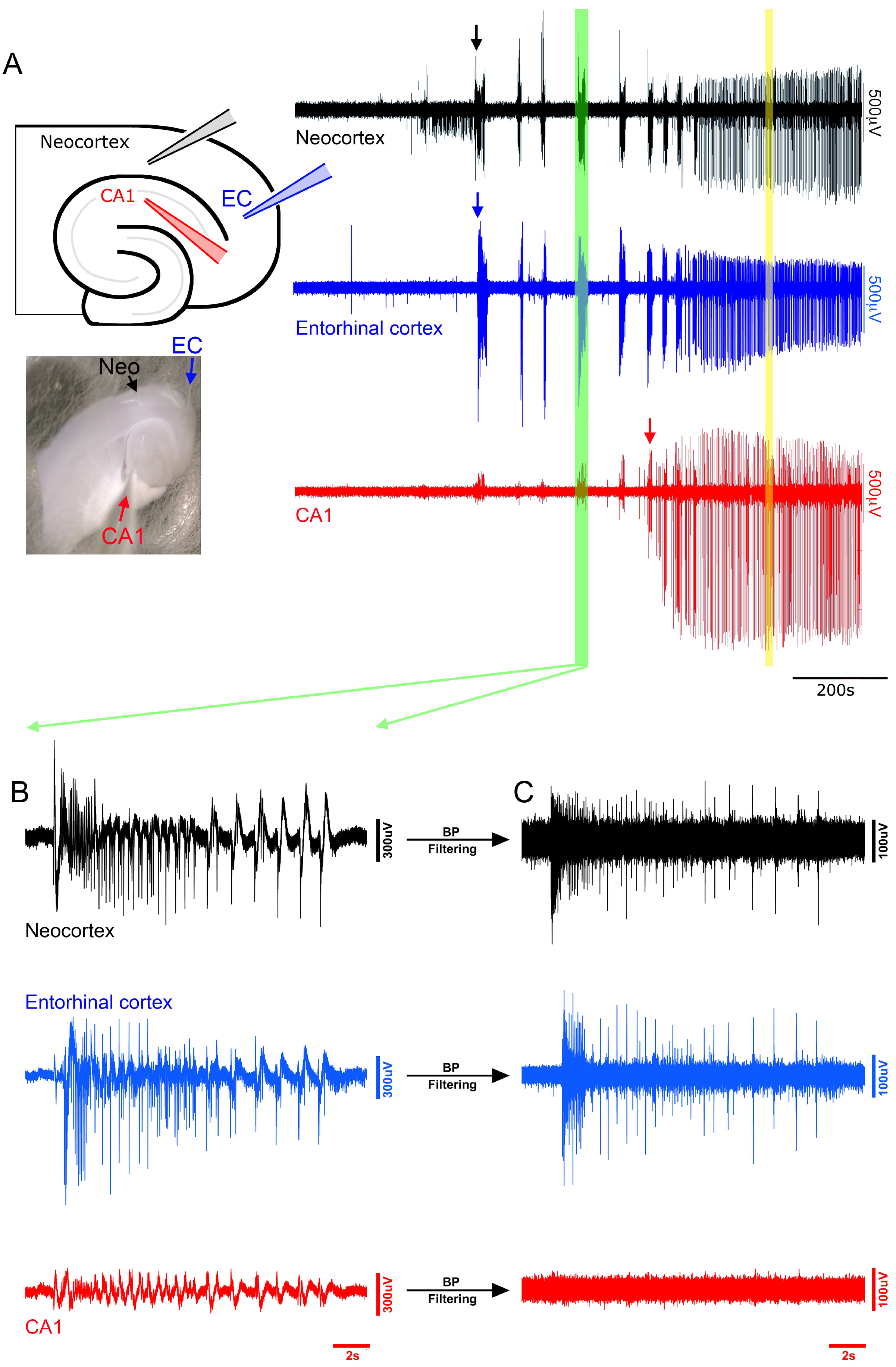
Typical pattern of evolving epileptiform activity following wash-out of Mg^2+^ ions from the bathing media (0 Mg^2+^ model), showing delayed recruitment of hippocampal circuits relative to neocortex. (A, left) Schematic and photomicrograph of a horizontal brain slice, showing the locations of three extracellular recording electrodes in hippocampus (CA1, red), the entorhinal cortex (EC, blue) and deep layers of the neocortex. (NC, black) and the corresponding, extracellular recordings (broad band), showing typical pattern of evolving epileptiform activity following washing out of Mg^2+^. The arrows indicate the first full ictal events, as indicated by intense multiunit (high frequency) activity, in the three recordings. The green vertical bar is expanded and represented in panels B and C. The yellow vertical bar is expanded and presented in Figure 2. (B) Broad band signals show small deflections in the hippocampal field at the time of large neocortical discharges, but high pass filtering (C) shows that these hippocampal signals are not associated with any significant unit activity.

**Figure 2.**
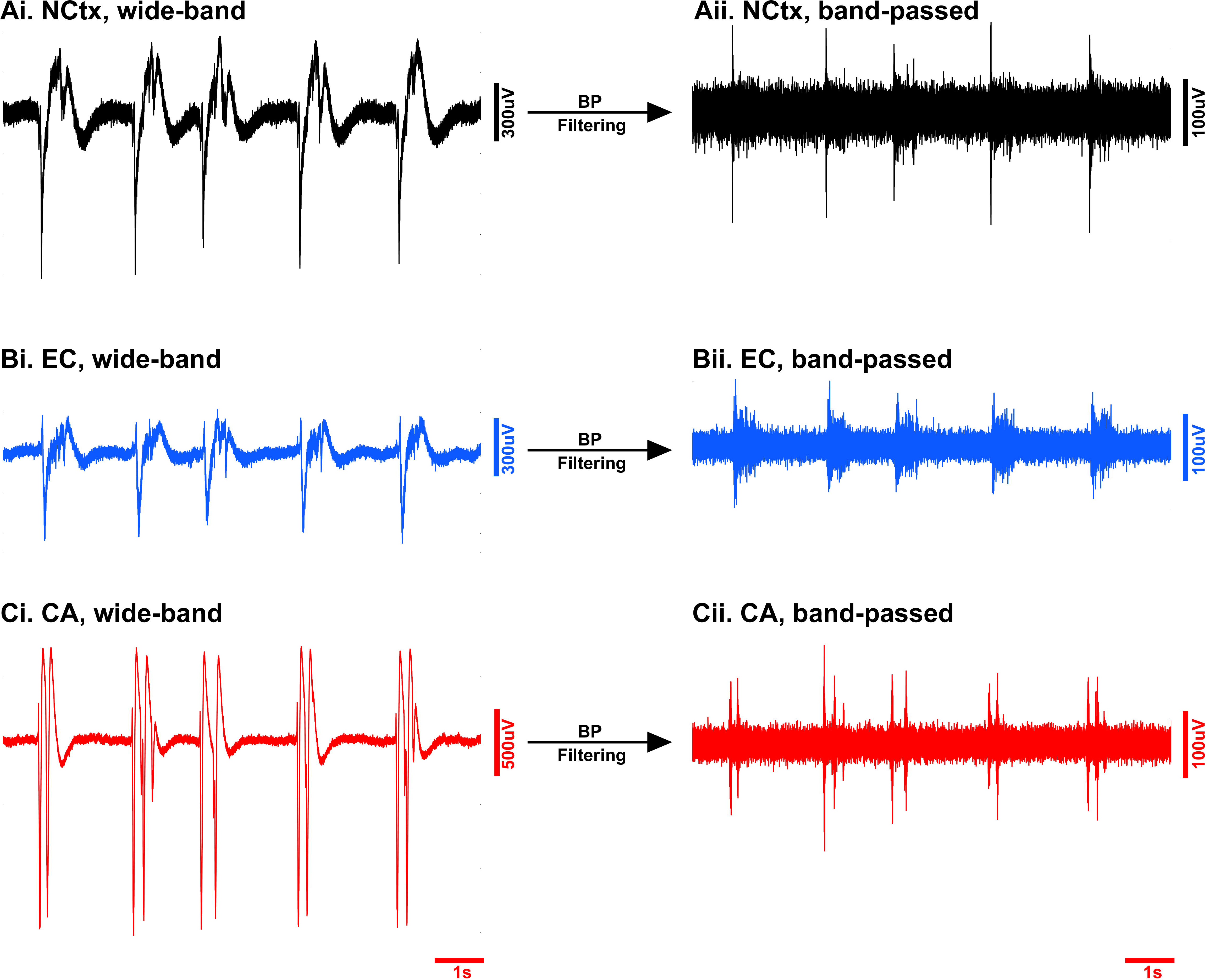
Late stage activity additionally involves hippocampal networks. Intense activity, involving a prominent high frequency component indicative of local network firing, is seen at all three electrode sites, in neocortex (A), entorhinal cortex (B), and for the first time also in hippocampus (C).

Recent studies of human extracellular recordings of epileptic discharges in humans have highlighted the importance of examining the high frequency component of epileptiform discharges to determine whether an event involves locally active neurons (Schevon *et al.*, 2012; Weiss *et al.*, 2013). In this regard, there appeared a striking difference between activity recorded in the hippocampus and the neocortical signals: the early events, including the tonic-clonic ictal events, were associated with only small field events in the hippocampus, and notably, with no measurable high frequency component (Figure 1A-C), indicating there is little local neuronal firing. We therefore considered these early events not to have invaded the local hippocampal networks. Using this high frequency component as the critical marker of ictal involvement, the first hippocampal ictal discharges occurred significantly later than the first neocortical discharges (Figure 1A arrows; Figure 3; Neocortex latency, 671 ± 41s; Entorhinal cortex, 699 ± 69s: Hippocampus, 2238 ± 284s; post-hoc Tukey test, p < 0.01). Epileptiform discharges in entorhinal cortex evolved in tandem with the neocortical discharges (Neocortex v Entorhinal, not significant; Entorhinal v Hippocampal, p < 0.01).

**Figure 3.**
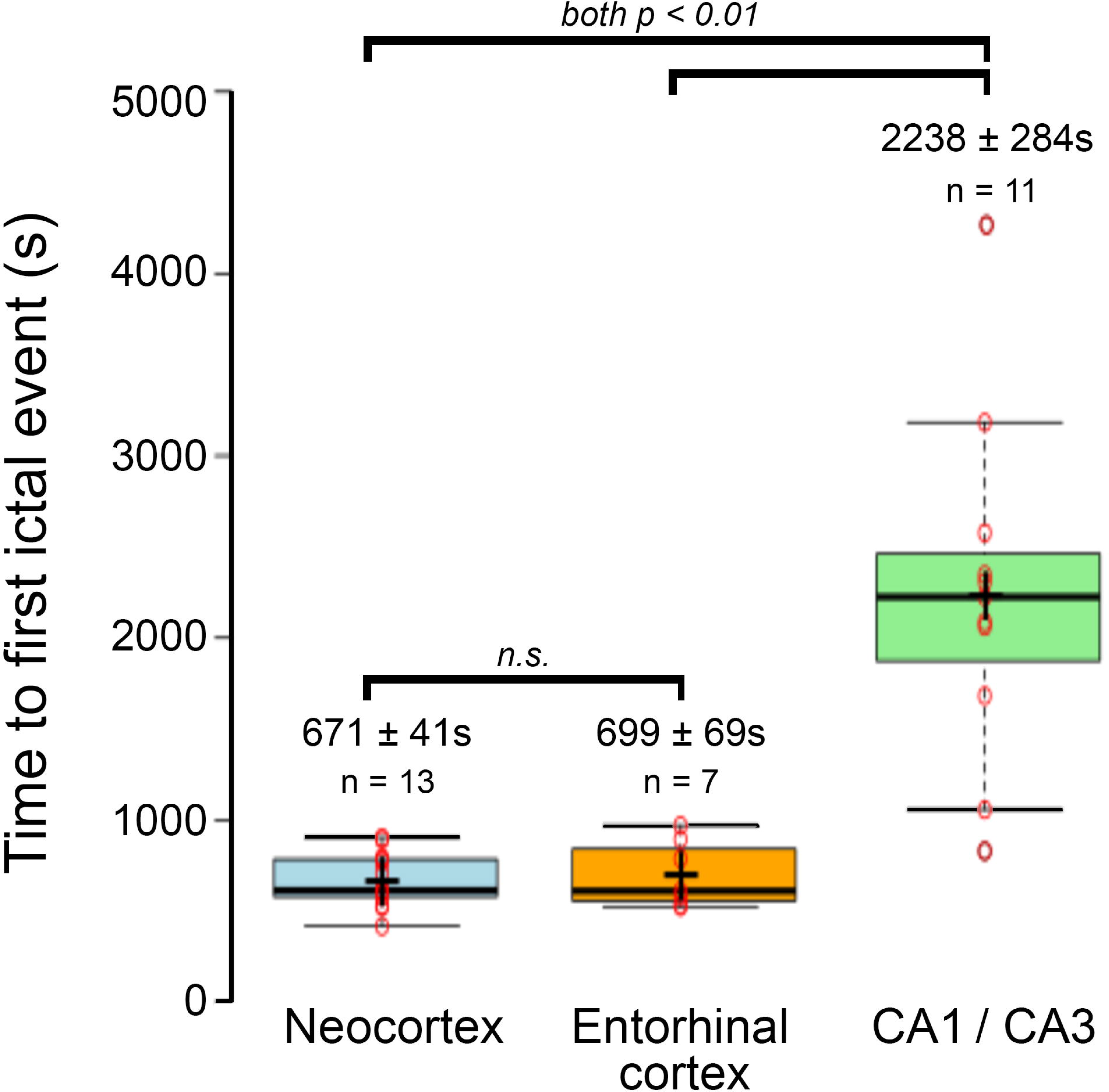
Ictal discharges are induced earlier in neocortical and entorhinal cortical networks than in hippocampus. Boxplot illustrating the pooled data, showing a highly significant delay of the earliest hippocampal epileptiform discharges relative to the first neocortical or entorhinal discharges (ANOVA F_[2,28]_ = 25.76, p < 0.001). The results of individual comparisons (post-hoc Tukey tests) are shown above the data distributions.

When finally, the hippocampal epileptiform discharges began, they showed a fundamentally different pattern, generally being a single large spike and wave discharge lasting up to 1.26 ± 0.11s (n = 10), or a short burst of discharges. In a further contrast to the prior neocortical activity, the inter-event intervals were short (2.98 ± 0.78s, n = 10), compared with the intervals between neocortical tonic-clonic ictal events (1^st^ – 2^nd^ event interval = 126.2 ± 17.2s; 2^nd^ – 3^rd^ interval = 117.1 ± 15.1s; 3^rd^ – 4^th^ interval =68.9 ± 10.6s). Interestingly, the pattern of neocortical discharges also changed once the hippocampal discharges started, to the same pattern of transient, but regular, spike and wave discharges. Discharges in the two structures, from this time forward, were tightly coordinated (Figure 2), but with the hippocampal discharges occurring before the neocortical unit activity (delay of onset of neocortical activity, relative to hippocampal activity = 87.1 ± 25.5ms, n = 8).

A noteworthy feature of these recordings was that tonic-clonic discharges appeared to be a hallmark only of neocortical recordings, with repeated events in every recording (n = 13). In contrast, we recorded such events in hippocampal electrodes in just 7.7% of the slices (1 in 13 recordings). There are, however, published records from rat brain slices of hippocampal tonic-clonic events ictal events (Swartzwelder *et al.*, 1987; Lewis *et al.*, 1989), but using much thicker brain slices (625μm). A key question then was whether this represented a species difference, or if instead, the thicker brain slices showed hippocampal tonic-clonic events because of better preserved neuronal connectivity. Our recordings were made typically from the middle sections in the dorsal-ventral axis, but we reasoned that on account of the curvature of the hippocampus, other levels may show different preservation of the axonal pathways. We therefore examined the more extreme slices, and discovered that the most ventral mouse brain slices (400μm, n = 4 slices) also showed tonic-clonic activity in CA3 (Figure 4A), as seen in thick (>600μm), rat sections; in this regard, therefore, there is no species difference. Notably, the tonic-clonic activity occurred in the entorhinal cortex before the CA3 region (Figure 4A), suggestive that the CA3 activity was conditional on the entorhinal cortex activity. This was confirmed by separating the two structures, after which the tonic-clonic pattern was maintained in entorhinal cortex, but abolished in CA3 (Figure 4B), which instead resorted to the spike and wave events already described. We conclude therefore that the early pattern of tonic-clonic activity is a hallmark of neo- and entorhinal cortex, and that instances of such activation in the hippocampus are downstream of activity at these other sites.

**Figure 4.**
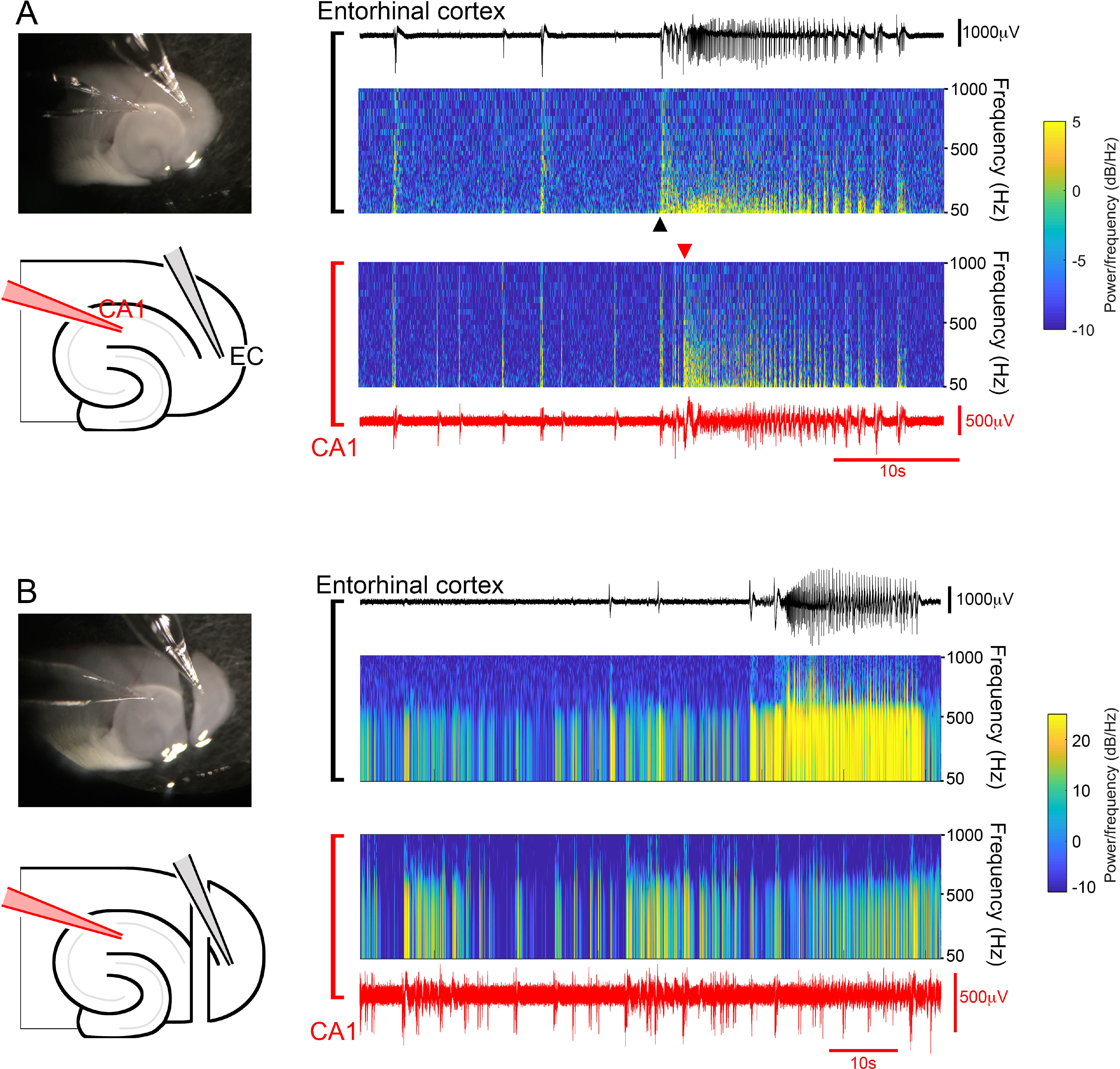
Tonic-clonic activation of CA1 can spread from entorhinal cortex. (A) Ventral horizontal brain slice, in which tonic-clonic activity spreads from entorhinal cortex into CA1. The spectrograms show that the sustained (tonic) high frequency component occurs first in the entorhinal cortex. (B) Recording from the same slice, following sectioning of the temporoammonic pathway. Note the tonic-clonic like event in entorhinal cortex, but not in CA1.

### Non-canonical propagation pattern of late epileptiform discharges

In contrast, during late stage activity, the hippocampal activity appears to be the pacemaker, entraining the other areas. We hypothesized that the entrainment is mediated through a polysynaptic pathway involving the entorhinal cortex. To test this, we cut away the caudal pole of the brain slice (we refer to these, henceforth, as “disconnected slices”), thereby entirely removing any potential synaptic pathway. Surprisingly, following the removal of the entorhinal pole, the hippocampal entrainment of neocortical discharges persisted unchanged (Neocortex, pre-cut rate = 0.47 ± 0.08Hz, post-cut = 0.47 ± 0.09Hz, n = 5, paired t-test, p = 0.96; Hippocampus, pre-cut rate = 0.48 ± 0.09Hz, post-cut = 0.49 ± 0.11Hz, n = 5, paired t-test, p = 0.83), and latency of onset of neocortical activity after hippocampal activity also remained unaltered (pre-cut = 71.1 ± 7.7ms, post-cut = 62.4 ±3.6ms, n = 5, paired t-test, p = 0.13).

In slices with the entorhinal pole removed from the start of the experiment (simultaneous with washing out Mg^2+^ ions), the evolving epileptiform activity showed the same general pattern in “intact slices” (slices including the entorhinal pole), albeit at a slightly slower rate (Figure 5A; Neocortex latency, 1035.71 ± 77.8s, n = 14; intact v disconnected, unpaired t-test, p = 0.0006). The first hippocampal ictal discharge occurred significantly later than the first neocortical discharge (hippocampus latency, 2356.10 ± 189.10s, n = 14; latency: hippocampus v neocortex, paired t-test, p = 0.0004; hippocampus latency, intact v disconnected, unpaired t-test, p = 0.79). And as with the intact slices, the start of the hippocampal discharges entrained the neocortical activity to the same pattern, despite the absence of any conventional polysynaptic connectivity between the two regions (Figures 5 and 6; neocortical latency, 57.8 ± 9.1ms; intact v dissected, unpaired t-test, p = 0.27). This entrainment was only lost when a second cut was made along the axis of the white matter bundle deep to the neocortical layer 6, thereby physically separating the neocortical and hippocampal networks (Figures 7). In these separate networks, the hippocampal discharge rate increased significantly (pre-cut, 0.35 ± 0.07Hz; post-cut, 0.49 ± 0.13Hz; n = 9, p = 0.0451; Figures 7C-E), whereas the neocortical discharge rate dropped significantly (pre-cut, 0.31 ± 0.05Hz; post-cut, 0.12 ± 0.02Hz; n = 9, p = 0.0099; Figures 7C-E). In tandem with the reduced rate of discharges in the neocortical networks, the duration of events showed a significant increase (pre-cut = 1.78 ±0.22s, post-cut = 6.86 ± 2.30s, n = 9, p = 0.048; Figure 7E). This result suggests that, in this late stage activity pattern, the interactions between hippocampal and neocortical networks are bidirectional: the hippocampal-to-neocortical influence is reflected in the pacing of neocortex by hippocampus; whereas the opposite influence is manifest as a mild brake on the hippocampal pacing, presumably by the tendency of the neocortical events to be extended, thereby also extending the refractoriness of the hippocampal pacemaker.

**Figure 5.**
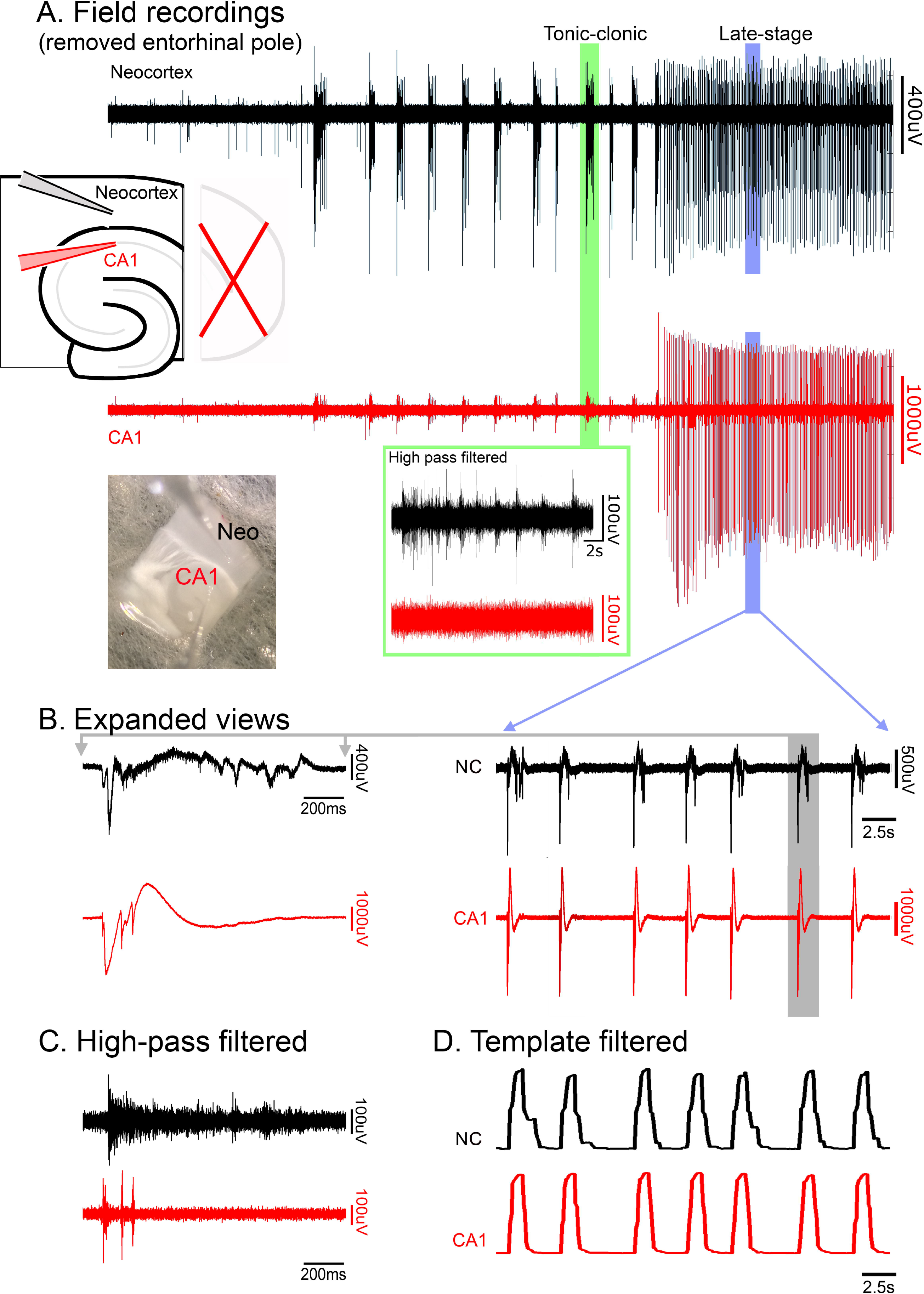
The late-stage epileptiform discharges are coordinated in hippocampal and neocortical networks through a non-synaptic pathway. (A) Extended recording of extracellular field potentials in CA1 and neocortex (NC), following wash-out of Mg^2+^, in a disconnected slice i.e., with entorhinal cortex removed, thereby disconnecting the two regions via any conventional multi-synaptic path. As in intact slices, the early discharges showed pronounced unit activity in neocortex, but not in the CA1 pyramidal layer (inset, green box). (B) Expanded view of wide-band, and (C) high-pass (>300Hz) filtered late stage activity in the same slice, showing prominent levels of unit activity in both territories. (D) The same traces filtered by a moving template of an average discharge.

**Figure 6.**
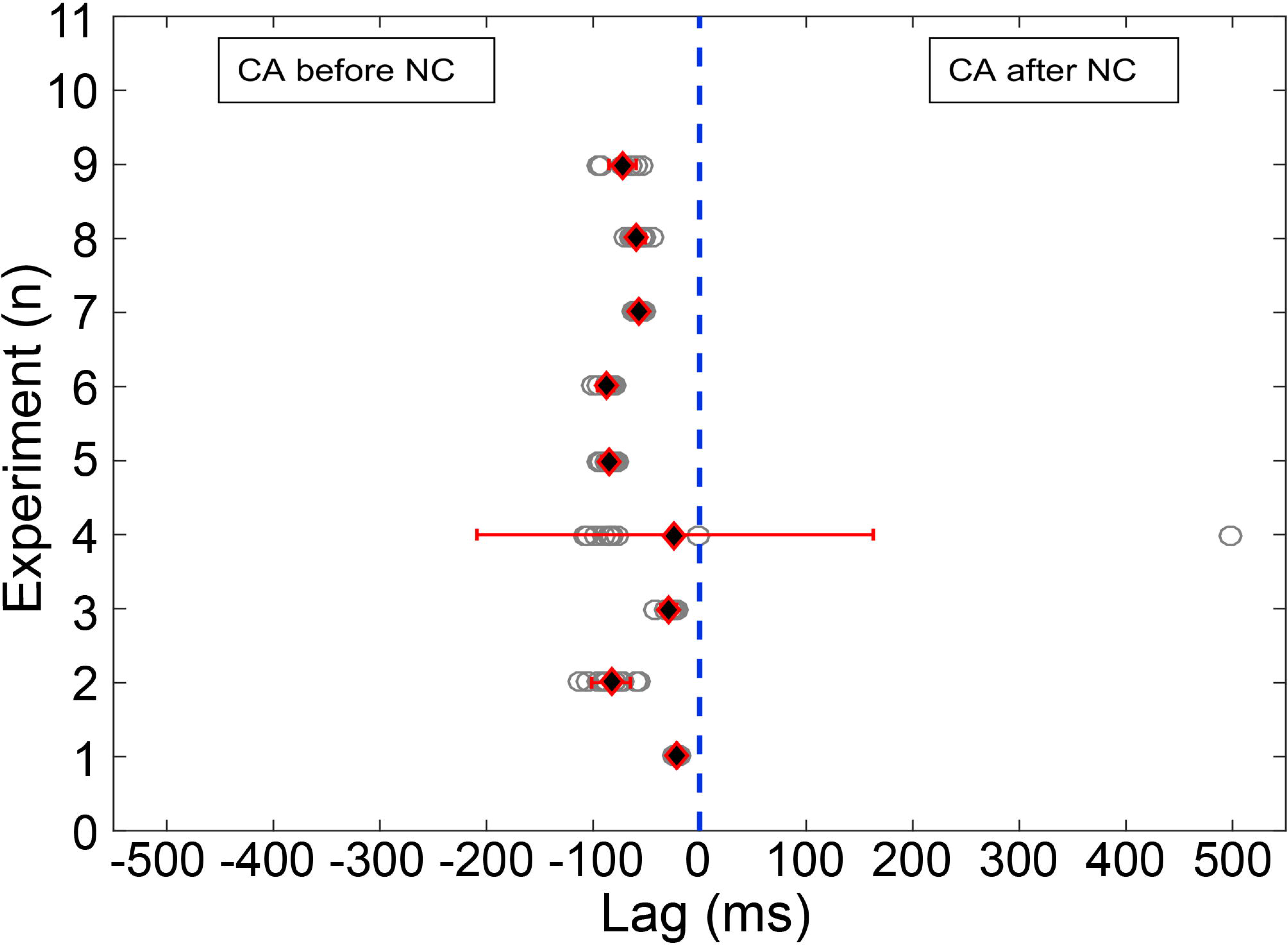
Hippocampal discharges precedes neocortical discharges during late-stage events. In synaptically-disconnected, hippocampal-neocortical slices, late-stage discharges in neocortex (NC) follow hippocampal (CA) discharges with a mean lag of 57.8±9.1ms (One-sample t-test: p<0.05, n=9).

**Figure 7.**
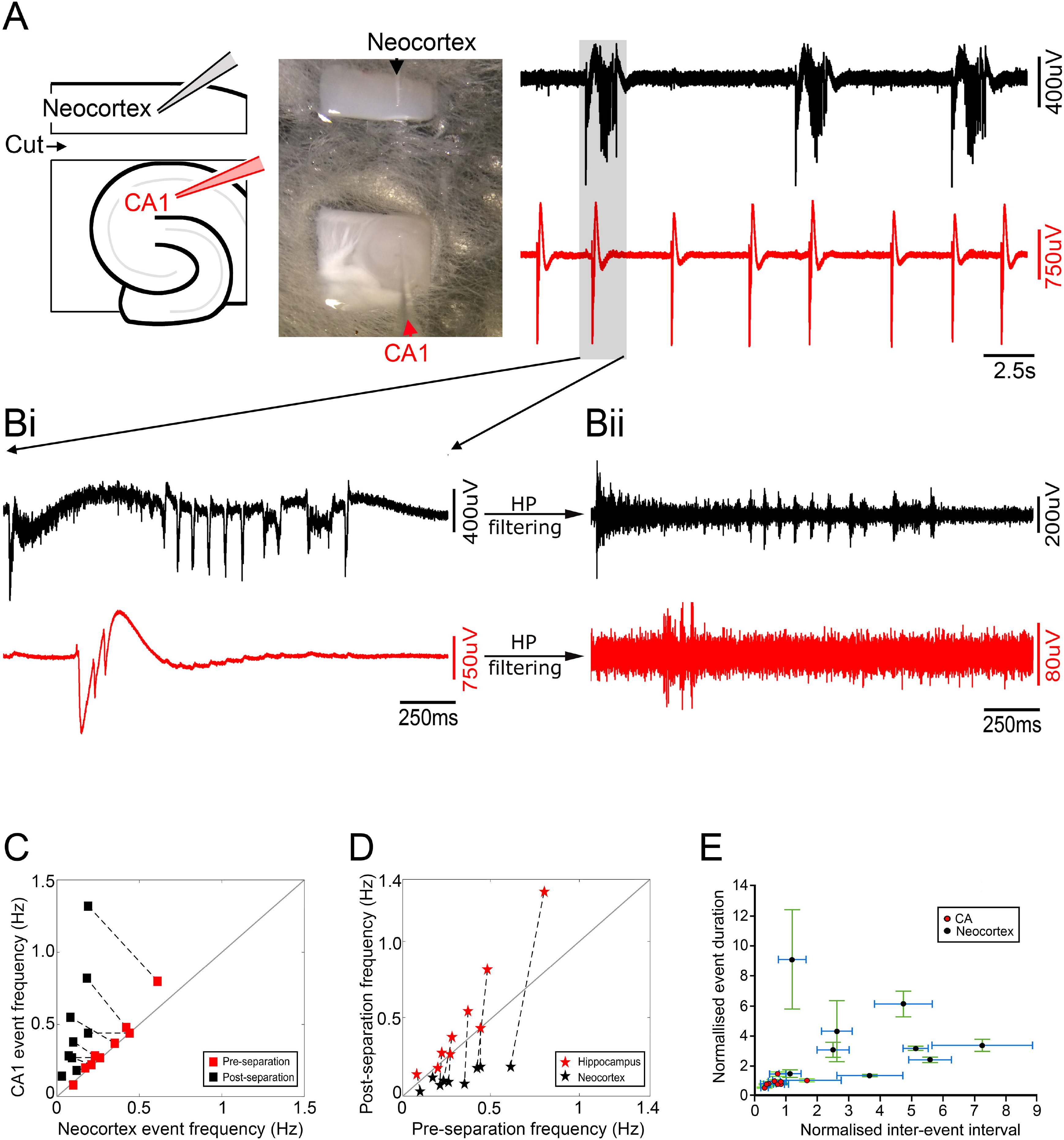
Entrainment of discharges is lost following physical separation of hippocampal and neocortical networks. (A) Photomicrograph and schematic showing the electrode placements in physically separated CA1 and neocortical areas, derived from a single horizontal brain slice, together with a period of late stage epileptiform discharges. Note the desynchronised discharges in the two territories, with a far slower rate of discharges in the neocortical tissue. (B) Further expansions show the broadband signal of the de-synchronised hippocampal and neocortical discharges. (Bi) Prominent unit activity is seen in both territories. (C) The relative rates of epileptiform discharges in the two territories before and after physically separated. In disconnected slices (“pre-separation”), the rates were equivalent (“pre-separation”, gray squares; CA: 0.35 ± 0.07 Hz, NCtx: 0.31 ± 0.05 Hz; n.s., n=9), but following physical separation of the tissues, the rates are significantly different (“post-separation”, black squares; CA: 0.49 ± 0.13 Hz, NCtx: 0.12 ± 0.02 Hz; p < 0.05, n=9). (D) Comparisons of discharge rates before and after physical separation of the hippocampal and neocortical tissues. Note how the neocortical data all fall below the line of unity, indicating a consistent slowing of the rate of discharges there (black stars; pre-separation: 0.31 ± 0.05 Hz, post-separation: 0.12 ± 0.02 Hz, p < 0.05, n=9). In contrast, the hippocampal data tend to lie above the line, indicative of an increase in hippocampal rate after the separation (gray stars; pre-separation: 0.35±0.07 Hz, post-separation: 0.49 ± 0.13 Hz, p < 0.05, n=9). (E) The duration and interevent intervals in the physically separated hippocampal and neocortical tissues, normalised to the values in the pre-cut brain slice.

Consistent with these opposite changes in rates, there was a highly significant drop in the correlation of events in the two networks (p = 4.1 × 10-7). We concluded from these experiments that the late stage epileptiform discharges arise in hippocampus, and these act as a pacemaker, driving discharges also in juxtaposed neocortical territories, and that this entrainment can occur independent of synaptic interactions (importantly, note that it does not exclude the possible involvement of conventional synaptic pathways in the development and propagation of epileptiform activity).

The interactions between the areas only really affected the late stage activity, because in slices that were dissected at the start of the experiment, to isolate the neocortex, entorhinal cortex and hippocampal subfields, the time to first ictal events in all three territories was unaltered, relative to recordings from intact slices (isolated neocortex, 607.3 ± 107.3s, n=6, unpaired t-test; vs intact: p=0.504; Entorhinal cortex, 1057.2 ± 200.1, n=5, unpaired t-test; vs intact: p=0.082; Hippocampus, 1595.4 ± 186.3, n=6, unpaired t-test; vs intact: p=0.1403).

One possible mechanism by which late-stage entrainment may happen is through diffusion of extracellular K^+^ from a local source of intense neuronal activation (Moody *et al.*, 1974; Heinemann & Lux, 1977; Somjen & Giacchino, 1985; Hablitz & Heinemann, 1987), thereby reducing the threshold for recruitment of other neighbouring territories. To assess the effects of [K^+^]_o_, rises, we made simultaneous recordings from 4 electrodes, at two sites, two located in neocortex and two in the CA1 region of the hippocampus, to record the local [K^+^]_o_, using an ionophore tip-filled electrode, and the local field potential. We found that the largest rises in [K^+^]_o_ associated with epileptiform events all occurred during the early tonic-clonic events in neocortex (Figure 8; note that much larger rises were, on occasions, recorded during spreading depression events, but these did not show high frequency activity denoting local neuronal firing). We analysed 24 events, in 6 brain slices, for which the multiunit activity showed that the event only occurred at one electrode site (Figure 8; 23 neocortical events, and 1 hippocampal discharge). Critically, in all cases, the site of unit activity was associated with a large rise in [K^+^]o (neocortical examples (n = 23), [K^+^]_o_ = 9.55 ± 2.70mM), but this did not spread to the other recording site (hippocampal [K^+^]_o_ = 3.57 ± 0.18mM; not significantly different from baseline [K^+^]_o_ = 3.50mM; Figure 8A, B). This was also the case for the single example of a prominent hippocampal discharge without neocortical involvement (hippocampal [K^+^]_o_ = 7.26mM; neocortical [K^+^]_o_ = 3.13mM). This showed that the [K^+^]_o_ changes are highly focal, indicating that this entrainment at a distance is not mediated by diffusion of K^+^.

**Figure 8.**
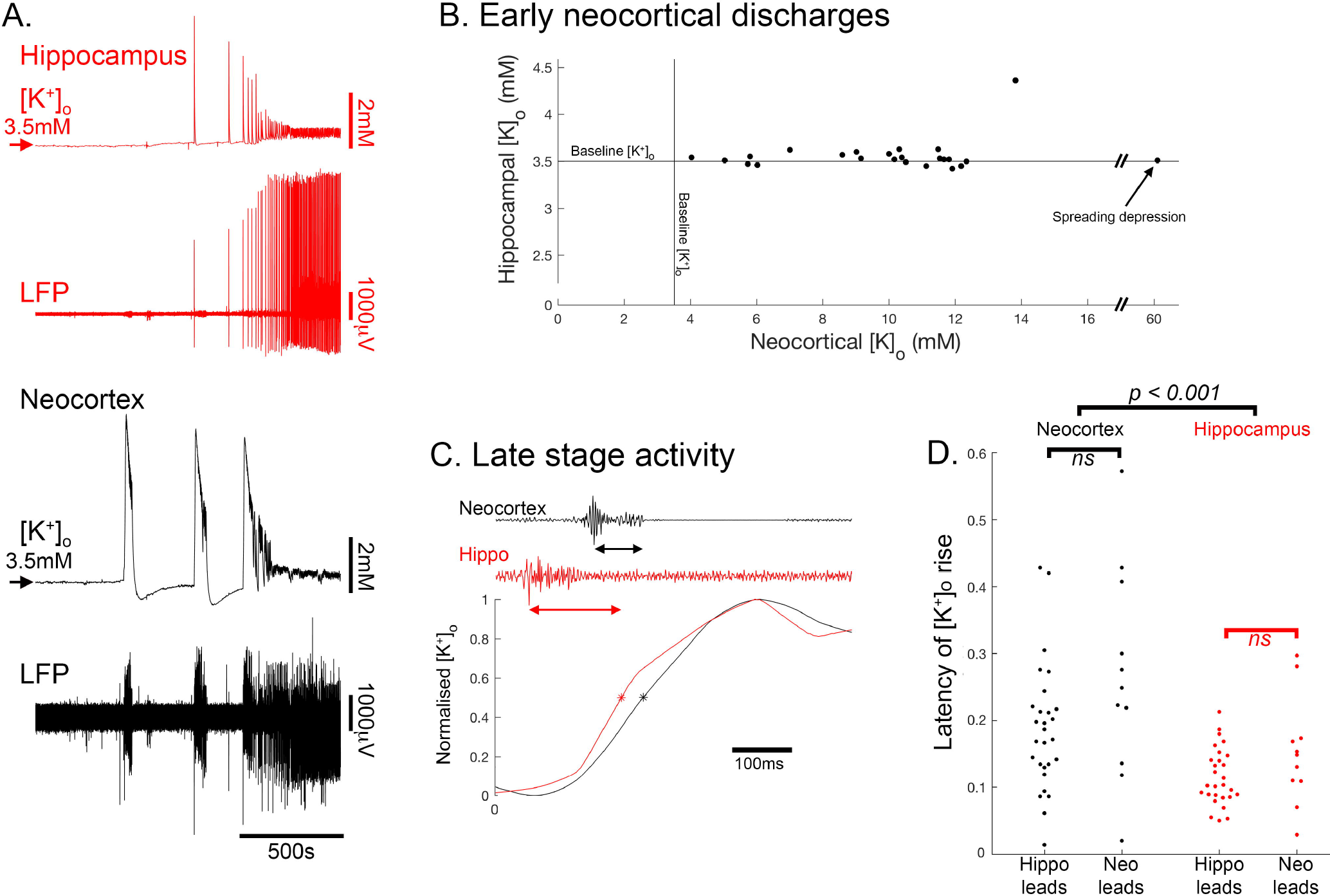
Entrainment does not happen through diffusion of K^+^. (A) Concurrent recordings of LFPs and [K^+^]_o_ in CA3 and neocortex. (B) Peak transient [K^+^]_o_, and the CA3 [K^+^]_o_ measures, during early neocortical tonic-clonic epileptiform events when there was no corresponding multiunit activity in CA3. (C) Latency measures during late stage events, with multiunit activity occurring in both neocortex and CA3 regions. (D) Population data, for the latency measures (41 data points from 6 brain slices) segregated by whether the neocortex or the hippocampal activity came first.

We next examined the late stage activity in which there was multiunit activation at both neocortical and hippocampal locations (41 events from 6 slices; Figure 8C, D). As expected, both sites also showed significant rises in [K^+^]_o_ associated with these bursts of local neuronal firing, and in all cases the main rise came after the local peak in the high frequency filtered LFP signal. There are, however, inherent problems with comparing timing between high and low band pass filtered signals, so we performed a further analysis comparing the latency of the rise in [K^+^]_o_ between events which led either in the hippocampal (n = 30) or neocortical (n = 11) territories. We reasoned that if activity in the follower territory was being triggered by a rise in [K^+^]_o_ diffused from the other site, then for those events, the rise would appear to occur significantly earlier relative to the local firing. In fact, there was no significant difference in latency between “leader” and “follower” events for either the hippocampal or neocortical recording sites (Figure 8D), leading us to conclude that in both groups, the local [K^+^]_o_ rise reflected, rather than caused, the local firing. We did observe that the latency for the [K^+^]_o_ rise in hippocampal circuits was significantly shorter than for neocortical circuits (p < 0.001), perhaps reflective of more densely packed neurons in the hippocampus. Collectively, these various analyses of [K^+^]_o_ rises associated with local neuronal firing, indicate that the entrainment of neocortical events by the hippocampal activity in this preparation does not happen by diffusion of K^+^ ions. Instead, it is likely to occur by the distant effects of a field potential onto circuits that are already highly excitable. This type of entrainment, we suggest, is also possible in vivo, during clinical epileptic events.

### Region-specific differences in drug sensitivity influences epileptic activity patterns

Previous work suggests that the different phases of evolving activity in this model activity may show differential sensitivity to drug manipulation. We investigated whether this may relate to regional sensitivity. One promising candidate is the GABA_B_ agonist, baclofen, which was reported to reverse the evolving pattern of activity, inducing a switch from what we term late stage activity (Swartzwelder *et al.*, 1987; Lewis *et al.*, 1989) (in the original description this was termed “interictal” activity; see Methods, Terminology) into tonic-clonic events. These experiments were performed on thick (600μm) rat brain slices, but since the activity generalized throughout the slice, the authors did not relate this to the source of activity. We repeated these experiments therefore on 400μm mouse brain slices to investigate whether the effect was location specific (Figure 9). We found that bath application of the GABAB agonist, baclofen (10μM), did indeed reverse the late-stage pattern (4 out of 4 slices), suppressing entirely the hippocampal bursting, and with the reappearance of tonic-clonic events in neocortex. Furthermore, if baclofen were applied from the start of the recording (when washing out Mg^2+^ ions), the tonic-clonic epileptiform events took longer to establish (0 Mg^2+^ latency = 609 ± 31s (n=5 slices); baclofen with 0 Mg^2+^ latency = 1573 ± 176s (n=4); p = 0.0005), but once that happened were maintained for the entire duration of the recordings, and hippocampal discharges never initiated. These results further support our conclusion that the different patterns of epileptiform bursting appear pathognomonic of the territories from which they originate.

**Figure 9.**
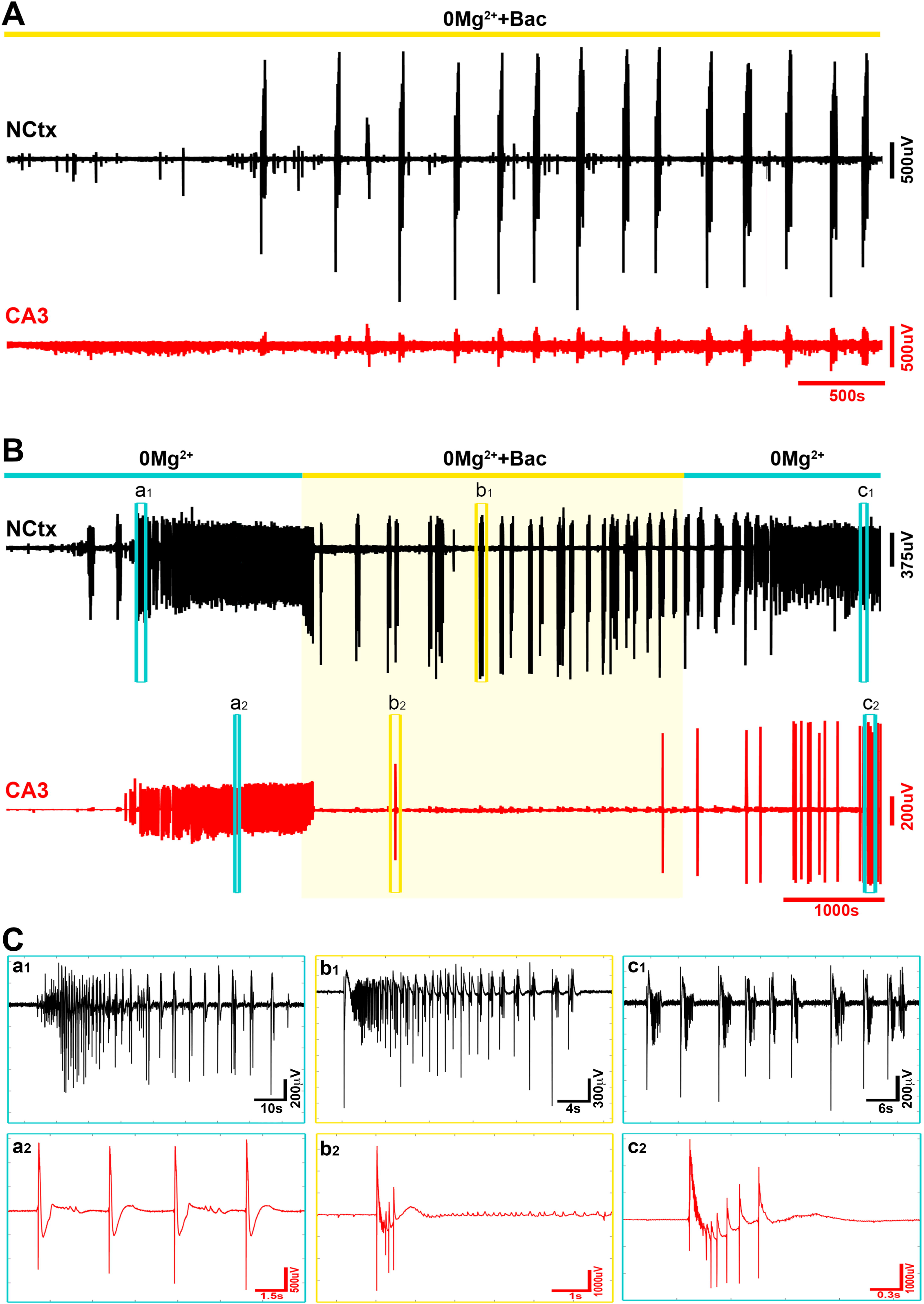
Hippocampal, but not neocortical epileptiform activity, is suppressed by GABAB activation. (A) The GABA_B_ agonist, baclofen, when applied simultaneous with the wash-out of Mg^2+^ ions, blocks any developing hippocampal activity, but does not suppress the development of tonic-clonic like events in neocortex. (B) Baclofen also blocks the hippocampal activity after it has started, thereby reversing the late stage pattern, and initiating the tonic-clonic like events that characterise the neocortical pattern. (C) Enlarged views of the discharges at the times indicated in B.

This region-specific difference also has a parallel in another under-appreciated feature of brain slice models, which is that application of the K^+^ channel blocker, 4-aminopyridine, has the exact opposite regional specificity to the 0 Mg^2+^ model, inducing epileptiform discharges in hippocampal territories significantly in advance of that in neocortex (Figure 10; neocortical latency = 605 ± 22s (n=9 slices); CA3 latency = 487 ± 17s (n=9); p = 0.04). Notably the early hippocampal activity in this instance, does not immediately entrain the neocortical territories, indicating that this entrainment requires changes in the local excitability.

**Figure 10.**
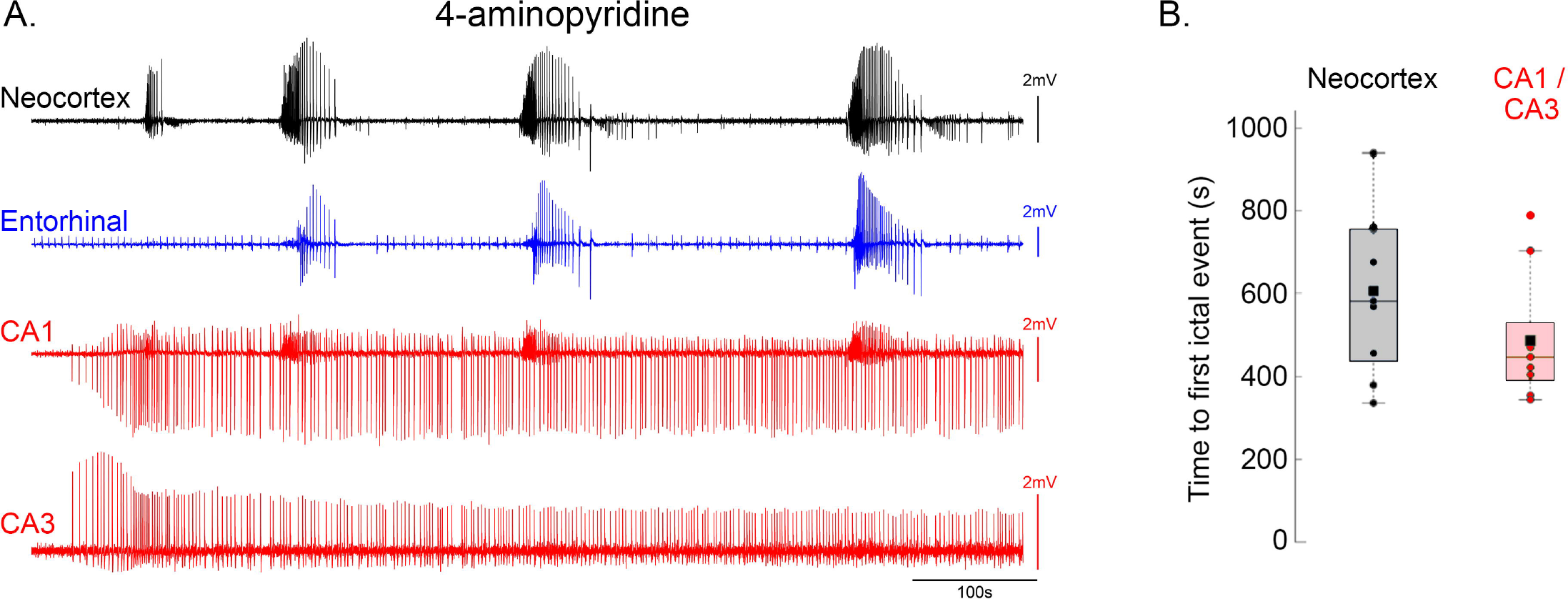
4-Aminopyridine activates hippocampal networks earlier than neocortical networks. (A) Quadruple electrode recording, showing the early activation of CA1 and CA3 in advance of neocortical epileptiform activation. Note that again the CA1 territory in this recording shows some tonic-clonic like activity that reflects the input from entorhinal cortex. (B) Pooled data showing that hippocampal discharges in this mode precede those in neocortex.

We conclude therefore that certain electrophysiological transitions arise from switches in the source of the pathological discharges, and reflect brain-region-specific differences in the propensity to support epileptiform discharges, and the electrophysiological signatures of these discharges. These switches between the focal sources can occur spontaneously, presumably reflecting local or cellular changes in the network excitability, but can also be influenced pharmacologically, indicating brain-region-specific differences, too, in their drug sensitivity.

## Discussion

We have provided demonstrations for several key principles of epileptic pathophysiology. The first is that transitions in the pattern of activity can reflect shifts in the source of discharges. A second key finding is that brain regions differ in how epileptic discharges manifest in the local circuits. A third principle is that when cortical networks have raised levels of excitability, they can be entrained through non-synaptic paths. It is important to realise that these are not the only changes underlying the development of epileptic activity (Whittington *et al.*, 1995; Fujiwara-Tsukamoto *et al.*, 2007; Ellender *et al.*, 2014). However, these various experiments do provide evidence of the interesting interplay between areas that are driving the pathology (the source, or ictal focus) and the susceptibility of secondary territories to be recruited. Of course, these acute brain slice preparations clearly do not incorporate all facets of the epileptic condition, but the substrates for all three of these principles do exist in vivo, and so, we would argue, all are likely to be relevant in spontaneously occurring seizures in humans.

The main focus of our studies was the marked transition from early tonic-clonic activity to a late-stage pattern of repeated, spike and wave discharges, occurring every 2-10s typically. This builds upon previous work done mainly using rat brain slices (Swartzwelder *et al.*, 1986b; Mody *et al.*, 1987; Anderson *et al.*, 1990; Dreier & Heinemann, 1990; 1991; Bragdon *et al.*, 1992; Morrisett *et al.*, 1993; Zhang *et al.*, 1995; Dreier *et al.*, 1998), but we extend this in two important ways. First is that, with the development of various mouse models carrying genetic mutations associated with human epileptic conditions (Yu *et al.*, 2006; Asinof *et al.*, 2015), our studies provide important confirmation that the evolving activity patterns in these models follow the same pattern in mice as they do in rats. This will facilitate a productive line of investigations regarding exactly how specific genetic mutations impact on network stability, using the 0 Mg^2+^ and 4-AP models to “stress-test” the tissue (Parrish & Trevelyan, 2018).

The second advance has been to clarify that different brain territories sustain characteristic epileptic discharge patterns. The extension of cortical area involvement with different pharmacological sensitivities, as well as the potential for non-canonical seizure propagation, may both contribute to pharmaco-resistance (Heinemann *et al.*, 1994a). The tonic-clonic pattern of discharges appears to be a feature only of discharges arising in neo-and entorhinal cortex, but not hippocampus; when such activity is seen in hippocampus, it appears to be relayed there from the entorhinal cortex. Note that we have not yet explored subicular and parasubicular territories. The hippocampal activity starts very late in the 0 Mg model, but very early in the 4-AP model. To the best of our knowledge, this key difference in these two very widely used models, which illustrates the principle about differing network susceptibility to seizures, has not been reported previously. The hippocampal discharges entrain the other territories in the late 0 Mg^2+^ model, but not in the early 4-AP activity, illustrating that the entrainment requires an increase in susceptibility (excitability) in the follower territories. Thus, while we emphasise that the explicit explanation of the transition is a shift in the source of the discharges, this must be underpinned by changes at the local network / cellular level, which alters the excitability of the networks.

Another important point, with clinical relevance, is that sudden changes in the local pattern of activity can be indicative of a shift in the source of the pathological driver. Previously we showed that a change in the direction of propagation of individual discharges is a marker of the passage of the ictal wavefront (Trevelyan *et al.*, 2007; Smith *et al.*, 2016). Our current study now shows that sudden changes in the pattern of activity recorded in neocortex reflects the appearance, or cessation (suppressed by GABA_B_), of a different pacemaker source, in this case within the hippocampal territories.

The late stage activity, which our studies indicate is a primarily hippocampal pattern, has been termed “interictal” activity by many observers, relating this to the clinical distinction between clinically manifest seizures, which presumably involve some motor territories in the brain, and epileptic electrophysiological discharges that are virtually clinically silent. The implication is that interictal discharges are restricted to areas that are less “eloquent”, but that disregards what may be more subtle effects on brain function. Indeed, increasing evidence exists now about the potential impact of interictal discharges on memory (Binnie *et al.*, 1987; Kleen *et al.*, 2010); such effects, in tandem without an explicit motor component, are entirely consistent with a hippocampal discharge. An important on-going clinical debate though has centred on the clinical significance of these events, specifically with regard to treatments predicated largely on the EEG findings. However, the clear demonstration that these are susceptible to GABA_B_ agonists provides a means to examine this. GABA_B_ agonists have been considered for treating epilepsy previously, but gave mixed results as assessed by seizure control (Terrence *et al.*, 1983). We suggest, however, that it might be considered as adjunctive therapy to more conventional anti-epileptics, with the aim of reducing interictal activity with a presumptive hippocampal origin, and thereby ameliorating memory dysfunction comorbidity. This is also consistent with recent work indicating that focal targeting of dentate function can impact on both memory issues and seizure severity (Liou *et al.*, 2018; Scharfman, 2018). Baclofen therapy is not entirely straightforward, because at different doses it appears to induce divergent effects on the hippocampus (Dugladze *et al.*, 2013), but it might be possible to calibrate the dose using EEG monitoring in individual patients.

Finally, we provide a proof of principle demonstration of entrainment of epileptiform discharges at a distance, through a non-synaptic mechanism. This is not mediated through diffusion of [K^+^]_o_, since any rises of [K^+^]_o_ appear to remain very local to the site of neuronal activity. Rather the entrainment is likely to arise through volume conduction of the field potential. Given the size of field fluctuations recorded even outside the skull during seizures, it is reasonable to presume that such entrainment across brain territories might also occur in spontaneous seizures, giving rise to complex patterns of spread. Of course, we stress that this demonstration of a non-canonical mode of spread does not downgrade the clear importance of conventional, synaptically mediated spread. A notable feature of this pattern of spread is that we only see it in a very particular situation, spreading into tissue that is already hyperexcitable, with a history of repeated epileptiform discharges. Thus, the specific instances of non-synaptic spread occur only in what we have termed “late-stage” epileptiform activity, in the 0 Mg^2+^ model; it does not occur with the early hippocampal discharges in 4-AP, nor in the early neocortical discharges in 0 Mg^2+^. However, the fact that separating the neocortex and hippocampus influences this late stage activity in both directions (the neocortex shows a significant slowing of the rate of discharges, whereas the hippocampal rate increases significantly) indicates that the interactions are indeed bilateral in hyperexcitable networks. This suggests first, that the neocortical discharges, which tend to last longer than the hippocampal ones, may impose additional refractoriness, and second, that the critical determinant of spread is that the follower network is “primed” for activation. We follow Jefferys’ nomenclature (Jefferys, 1995) in avoiding the use of the term “ephaptic spread”, since he reserves this term for activation of juxtaposing cells (it derives from the Greek word “to touch”), whereas the effect we describe clearly occurs at a distance. We suggest that this occurs through a distant field effect. Even though this might be considered a relatively weak effect, there is, however, an important precedent of this result, whereby epileptiform discharges can be entrained by minimal activation in an already hyperexcitable network: this is the demonstration that bursts of action potentials of a single pyramidal cell can entrain these discharges in disinhibited hippocampal networks (Miles & Wong, 1983). In conclusion, we have demonstrated several key principles of network interactions in epileptic pathophysiology. Although the precise nature in which they will be manifest may be slightly different in a chronically epileptic subject, these phenomena are highly likely to be relevant also in vivo, and may inform our interpretation of clinical electrophysiology.

## Acknowledgements

We’d like to thank Dr Yujiang Wang and Prof John Jefferys for comments on the initial write-up of this project, and to Prof Juha Voipio for help setting up the [K^+^] measurements.

